# Protein Language Visualizer: A Repository for Homology Exploration with Language Model Embeddings

**DOI:** 10.1101/2024.11.19.624229

**Authors:** Javier Espinoza-Herrera, María F. Manríquez-García, Sofía Medina-Bermejo, Ailyn López-Jasso, Juan Pablo Ruiz-Alcocer, Adriana Siordia, Sarah M. Veskimägi, Nathaniel Roethler, Adrian Jinich

## Abstract

The era of modern AI-driven representations of proteins is here, and moving fast, yet tools for their intuitive visualization and exploration lag behind. Sequence Similarity Networks (SSNs) have long filled this role for alignment-based methods, providing simple but widely adopted platforms for grouping proteins by homology. Building on this foundation, we present the Protein Language Visualizer (PLVis), a modular framework that applies existing pre-trained protein language model (pLM) embeddings, dimensionality reduction, and clustering to generate interactive maps of protein relationships. The central contribution is the PLVis Repository, an online resource where thousands of reference proteomes can be compared and annotated through an accessible, interactive interface, much like SSNs became impactful not for their technical novelty but for their broad usability. We first validate that well-separated clusters in PLVis reliably capture homology information, while emphasizing caution when interpreting central “fuzzy” regions. We then illustrate the value of PLVis through case studies spanning individual protein families to full proteome comparisons across Mycobacterium and Plasmodium species. By combining methodological clarity with broad accessibility, the PLVis Repository provides a low-barrier platform for exploring proteomes through the lens of language models.

## 1 Introduction

High-throughput sequencing has accelerated protein discovery, far outpacing functional annotation [1]. UniProtKB now contains over 250 million sequences, yet fewer than 1% are in Swiss-Prot, the manually reviewed section [2]. Even with automated pipelines, the function of more than 30% of protein-coding genes remains unknown [3, 4].

Especially in the new era of AI-driven protein representations, where models are powerful but often black-box, visual tools are invaluable for exploring and interpreting large protein collections. Sequence Similarity Networks (SSNs) have long served this role and remain widely adopted [5, 6, 7, 8, 9]. SSNs provide a simple yet effective way to display protein relationships: nodes represent proteins, and edges reflect pairwise similarity scores, which may be scaled or filtered according to the underlying metric [10]. Tools such as CLANS have long supported the visualization of pairwise sequence similarities, further contributing to the widespread use of network-based representations [11]. More recently, web tools like the EFI-EST have made SSNs accessible and popular for probing sequence-function relationships [12].

Other statistical approaches, such as Profile Hidden Markov Models (HMMs), are also widely used to detect conserved patterns in protein sequences and classify them into families and domains [13, 14]. Originally developed in speech recognition and later adopted in NLP [15, 16], HMMs underpin many of the curated resources biologists rely on today. However, unlike SSNs, HMM-based results are not typically designed for direct interactive visualization; instead, they are represented through sequence logos, hierarchical trees, or heat maps [17, 18, 19]. Although conceptually distinct, the similarity scores produced by HMM profile searches can also be used in network-based analyses when desired. Complementing these sequence-based strategies, large-scale analyses of structure databases (e.g. AlphaFold Protein Structure Database) highlight how structure-based comparisons can also accelerate the annotation of uncharacterized proteins [20].

Protein Language Models (pLMs), also adapted from NLP, are the conceptual successors of HMMs [21, 22]. Built on transformer architectures with attention and positional encoding [23, 24], they are trained through masked-sequence prediction to learn context-dependent representations of amino acids. The resulting high-dimensional embeddings can represent both individual residues (tokens) and entire protein sequences, providing powerful features for downstream prediction and classification tasks, including structure prediction, as well as protein generation and design [23, 25, 26, 27].

The rise of pLMs and their rich embeddings presents an opportunity to design a new generation of interactive visualization tools for protein similarity. Here, we highlight an accessible approach: interactive exploration of two-dimensional projections of pLM embeddings. Beyond the method itself, our aim is to build a shared resource where such visualizations can be systematically applied across proteomes and protein families. By turning what could be one-off plots into a collective, searchable repository, we hope to make these representations broadly useful for researchers, educators, and even community-driven or citizen science efforts. While several studies have combined protein-language model embeddings with dimensionality reduction or embedding-space visualization to analyze individual protein families or functional subsets (e.g., via embedding trees or interactive embedding-space exploration), these efforts remain largely family-centric and do not systematically address species-wide proteome comparison [28, 29, 30, 31]. To our knowledge, no existing resource provides a taxonomically organized, interactive repository for embedding-based visualization across full reference proteomes, integrating clustering, annotation enrichment and cross-species comparative capability.

In this work, we focus on how protein language model embeddings can be explored through 2D projections. Aware of critiques of dimensionality reduction in other fields, particularly single-cell genomics [32, 33, 34, 35, 36], we cautiously assess how well these projections preserve sequence relationships. We find that well-separated clusters consistently retain homology information, whereas large, central “fuzzy” regions require caution. Next, through case studies, we illustrate how such projections can support exploratory analysis across scales, from individual protein families to full proteomes, highlighting examples from *Mycobacterium* and *Plasmodium* species. Finally, we introduce the PLVis Repository, an interactive resource covering thousands of reference proteomes, alongside a Colab notebook that enables researchers to apply the pipeline to their own data. Together, these contributions establish the PLVis Repository as a low-barrier platform for comparative proteomics in the era of protein language models. As with SSNs, the strength of PLVis lies not in methodological novelty but in accessibility and scale: by turning language model embeddings into reusable visual resources, we aim to make them as widely useful for the protein community as SSNs have been for sequence alignments.

## 2 Results

### 2.1 Evaluating PLVis projections: methodological choices, SSN comparison, and functional enrichment

We begin by outlining the PLVis pipeline (Figure 1). Protein sequences are first embedded with a pLM (e.g., ESM2, ProtT5), then reduced to two dimensions with algorithms such as UMAP, t-SNE, or TriMAP. Clustering methods (e.g., K-Means, DBSCAN) identify groups of related proteins, which can be automatically labeled by frequent terms in their annotations. The pipeline is modular, allowing users to mix and match embedding models, dimensionality reduction, and clustering approaches [37, 38, 39, 40, 41]. In the following sections, we use this framework to show how PLVis captures patterns consistent with established homology and orthology annotations, complementing sequence similarity networks and enabling comparative analyses across entire proteomes.

**Figure 1:**
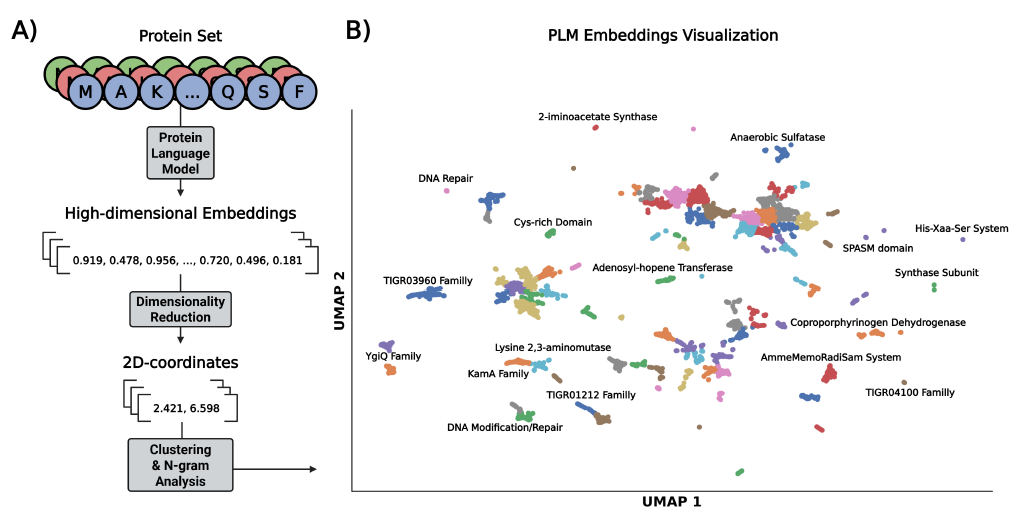
Schematic overview of the PLVis pipeline. (A) A set of protein sequences is fed to a PLM to obtain embeddings. These embeddings are reduced to two dimensions with a dimensionality reduction algorithm. Next, each point is clustered with its neighbors, and a title that averages the most common words in the protein names is generated using bi-gram analysis. (B) Example visualization of the processed data. The arrows indicate the flow of information through the pipeline, where each step can be performed using the model and reduction algorithm of choice.

For the analyses in this study, we drew on five datasets: 10,000 radical SAM (rSAM) enzymes, a set of sterol-binding proteins, the full proteome of *M. tuberculosis*, eight *Mycobacterium* proteomes, and five *Plasmodium* proteomes. To compare dimensionality reduction methods, we generated projections with UMAP, t-SNE, and TriMAP under default parameters (Figs. S1-S2) and assessed clustering quality using the Davies-Bouldin Index (DBI) and Calinski-Harabasz Index (CHI). For the smaller datasets, UMAP consistently produced more compact and well-separated clusters, while for proteome-scale datasets, its performance was intermediate. We further examined UMAP hyperparameters (e.g., min_dist, random seeds) and found that the overall clustering patterns and case study conclusions remained robust (Fig. S3). Given this balance of performance and interpretability, we selected UMAP as the default method for subsequent analyses. Lastly, because PLVis is designed as a visualization and exploration tool, clustering is applied to the two-dimensional projections rather than to the original embedding space, ensuring that cluster boundaries correspond to visually separable regions in the final map; this distinction between high-dimensional neighbors and 2D cluster structure is illustrated in Fig. S4. For consistency across datasets, the number of clusters (K) was selected by maximizing the silhouette score under default parameters, providing an automated way to identify visually coherent groups.

Next, we compared PLVis to the standard BLAST-based approach, the Sequence Similarity Network (SSN). SSNs are widely used to explore sequence–function relationships by grouping proteins into connected clusters based on pairwise alignment scores [9, 42, 43, 44]. A key feature of SSNs is their reliance on a user-defined threshold: at high cutoffs, many proteins appear as isolated nodes, whereas at lower cutoffs more connections are drawn but functional specificity may be lost. To examine how this thresholding compares with PLVis, we analyzed both the 10,000 randomly selected radical SAM (rSAM) enzymes and sterol-binding protein datasets using both approaches (Fig.2).

Figure 2A1-B1 shows that the embeddings of both datasets form distinct clusters in two-dimensional space. We selected the five densest clusters from the SSNs (Figure 2A2-B2) and examined their correspondence within the PLVis projections, which were partitioned using K-means clustering, yielding 88 clusters for the rSAM dataset and 48 clusters for the sterol dataset. Overall, SSN clusters were largely conserved within the PLVis projections. For instance, SSN cluster 4 in the rSAM comparison was fully contained within PLVis clusters 7 and 57, while SSN clusters 3 and 5 in the sterol-binding comparison were each confined to a single K-means cluster (21 and 32, respectively). A salient characteristic of the SSNs is the large fraction of proteins that appear as isolated nodes at the selected similarity threshold, comprising 1,932 proteins ( ∼ 20% of the rSAM dataset) and 4,420 proteins ( ∼ 80% of the sterol dataset). In contrast, within the PLVis representations, these proteins were redistributed across 75 of the 88 K-means clusters in the rSAM dataset and across all clusters in the sterol dataset, thereby integrating them with related sequences rather than leaving them disconnected.

**Figure 2:**
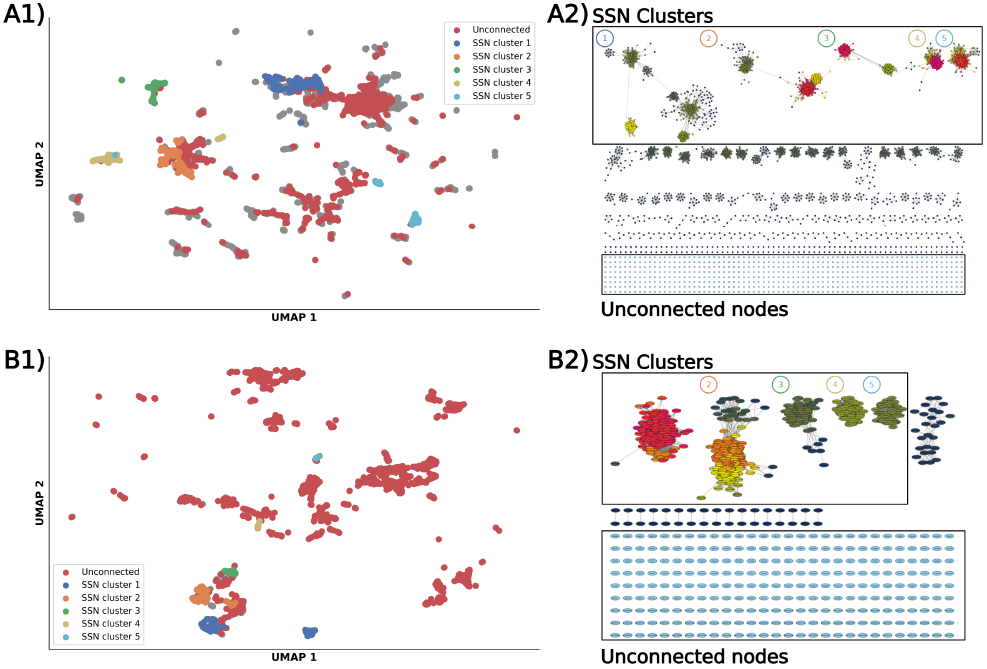
Comparison of PLVis and corresponding sections of the SSN for the (A) rSAM and (B) sterol datasets. (1) Each color in the PLVis represents a cluster in the SSN (blue:1, orange:2, green:3, yellow:4, cyan:5). Proteins colored red in the PLVis represent proteins that are unconnected in the SSN. (2)The SSNs were generated using pairwise sequence alignment scores selected to approximate a 35% sequence identity cutoff. The 5 densest clusters were selected and named accordingly. Nodes situated inside the unconnected region represent proteins that were cut from the threshold.

**Figure 3:**
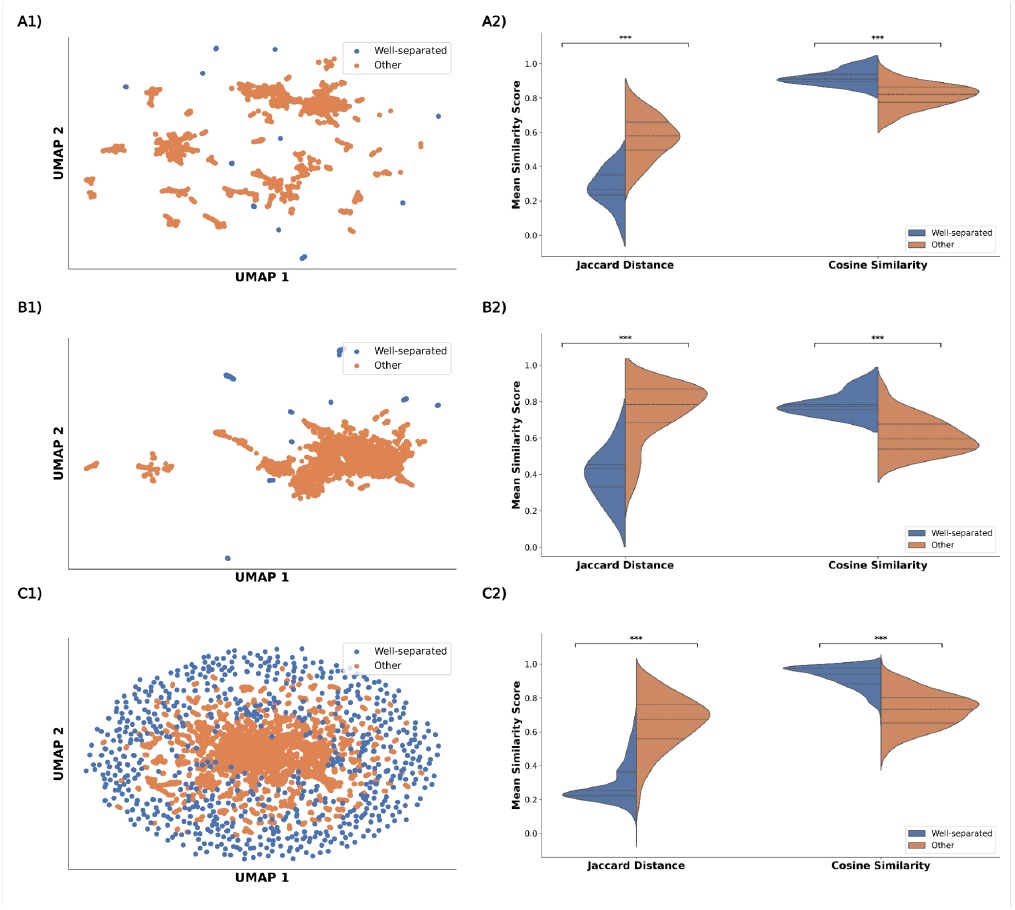
Well-separated clusters of data are statistically better at conserving high-dimensional data. (1) UMAP plots of PLM embeddings for (A) 10,000 radical SAM enzymes, (B) *M. tuberculosis proteome*, (C) 8 *Mycobacterium* genus proteomes; blue - well-separated clusters, detected by silhouette score above threshold (S ≥ 0.95), orange - clusters with a silhouette score below the threshold. (2) Violin plots of the average Jaccard distance of proteins and cosine similarity of high-dimensional embeddings within well-separated clusters (blue) and the rest of the clusters (orange). Statistical comparison was performed using the Mann-Whitney U test (***p < 0.001).

To evaluate the extent to which PLVis clusters capture biologically meaningful information, we performed enrichment analyses on all K-means clusters using a hypergeometric test for both datasets. For each cluster, the most frequent InterPro annotations (Family, Domain, and Other) were assessed relative to their background frequencies in the full dataset. Among clusters with available annotations, 96% were enriched for an InterPro “Family” term, 74% for “Domain,” and 68% for “Other” in the rSAM dataset, while corresponding values for the sterol dataset were 95%, 90%, and 100%, respectively. Importantly, 93% of the 1,932 proteins that appeared as isolated nodes in the rSAM SSN were assigned to PLVis clusters enriched for an InterPro “Family” annotation. For example, PLVis cluster 45 grouped 18 previously unconnected proteins with sequences annotated as belonging to the TatD-associated rSAM family (IPR023821). Likewise, in the sterol-binding representation, PLVis cluster 16 grouped 143 SSN isolated proteins that share a conserved site (IPR020904) within the short-chain dehydrogenase/reductase family (IPR002347). These results illustrate how PLVis can recover meaningful groupings for proteins that SSNs leave disconnected and often excluded from downstream analyses.

To further assess whether PLVis clusters align with curated homology and orthology classifications, we performed systematic enrichment analyses against CATH FunFams and OrthoDB ortholog groups across all datasets (Fig. S5). For each k-means cluster, enrichment was evaluated using a hypergeometric test with Benjamini–Hochberg correction. OrthoDB annotations showed widespread enrichment across clusters in all datasets, indicating strong consistency between embedding-derived groupings and curated ortholog assignments. In contrast, enrichment for CATH FunFams was more variable, reflecting the more limited coverage of structural annotations for large and diverse enzyme families such as rSAM. Together, these results support the use of PLVis as an exploratory framework for organizing proteins in a manner consistent with established homology and orthology resources, without performing explicit homology or orthology inference; however, the resulting organization is contingent on the underlying embedding model used to represent protein sequences.

### 2.2 PLVis projections preserve local information within well-separated clusters

Aware of the well-documented shortcomings of 2D projections in other biological contexts, we analyzed the properties of PLM embedding projections. In single-cell sequencing studies, dimensionality reduction methods such as t-SNE and UMAP often distort high-dimensional structure. This has been quantified using the Jaccard distance, which measures the overlap between the N nearest neighbors of each point in the original high-dimensional space (ambient space) and in its 2D projection (embedding space). Across 14 single-cell datasets, average Jaccard distances of ∼ 0.7 were reported, indicating substantial loss of local neighborhood information [32, 36]. Such distortions can be considered in terms of local preservation (relationships within clusters), and global preservation (the arrangement of clusters relative to each other in 2D).

We applied this same metric to pLM-based projections to quantify how much clusters preserve neighborhood information. Guided by interactive exploration of annotations across clusters, we hypothesized that well-separated clusters, compact groups that are visually distinct from large, central “fuzzy” regions, more faithfully preserve local relationships in the ambient embedding space, and that this distinction could be quantified rather than assumed. To evaluate this, we analyzed five UMAP datasets, whose properties are summarized in Table S1.

To quantify cluster separation, we calculated silhouette scores for each protein at a fixed number of clusters. We tested thresholds from 0.5 to 0.95, and found that higher cutoffs consistently enriched for clusters with stronger agreement between 2D projections and embedding-space similarity. For clarity, we used 0.95 to illustrate the most stringent case, but the same trend was observed across thresholds (Figs. S6) (shown in blue in Fig.3). For each protein, we then calculated the Jaccard distance between its neighbors in the ambient embedding space and in the 2D projection. In our analysis (Figs.3, S7), well-separated clusters had significantly lower Jaccard distances than other clusters (p < 0.001, Mann–Whitney U). As a complementary measure, we compared cosine similarity of high-dimensional embeddings within clusters, which was also significantly higher for well-separated clusters (p < 0.001), reflecting that they contain more closely related proteins.

Among the datasets, the eight *Mycobacterium* proteomes showed the clearest effect, with the highest number of well-separated clusters and the strongest agreement between 2D projections and embedding-space similarity. This likely reflects the presence of many conserved protein families shared across closely related species, which are expected to form tight and well-defined groups in embedding space. Importantly, this pattern emerges most clearly in cross-species comparisons, where proteins annotated to the same conserved families cluster together and separate from lineage-specific proteins. Such comparative contexts make it particularly straightforward to visually and quantitatively explore conserved protein families across proteomes. Although these trends are expected given the properties of non-linear projections, explicitly quantifying them provides practical guidance for interpreting PLVis visualizations, clarifying when visual separation reflects meaningful embedding-space structure and when it does not.

We then sought to validate the well-established fact that inter-cluster distances in non-linear projections are not particularly meaningful by evaluating whether nearest neighboring clusters have more similar pLM embeddings than randomly selected clusters. Given that non-linear dimensionality reduction techniques like t-SNE and UMAP warp the shape of the data when projecting to lower dimensions, distances between clusters of data points should not be interpreted directly. Using the previously mentioned datasets, we calculated the average cosine similarity for the embeddings of proteins within each cluster and compared it to the inter-cluster cosine similarity with (1) the nearest neighboring cluster and (2) a randomly selected cluster (Figure 4 & S8).

**Figure 4:**
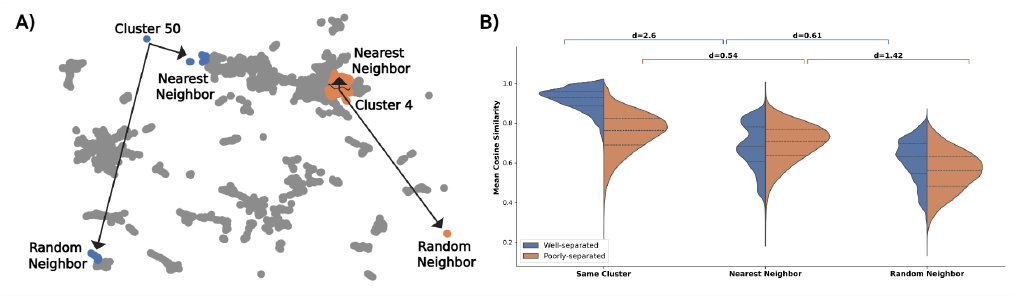
Distance between clusters in the PLVis projection is not associated with sequence similarity. (A) Example schematic of a well-separated cluster (blue) and a poorly-separated cluster (orange) and their relative positions to their corresponding closest neighboring cluster and a randomly selected cluster; well-separated clusters, detected by silhouette score above threshold (S ≥ 0.95), poorly-separated clusters with a silhouette score below threshold (S < 0.5). (B, C, D) Violin plots of the mean sequence similarity score for each cluster when comparing its proteins with the nearest neighboring cluster and a randomly selected cluster for (B) 10,000 rSAM enzymes, (C) *M. tuberculosis* proteome, (D) 8 *Mycobacterium* genus proteomes. Significance bars represent the effect size between sets using Cohen’s D.

In Figs. 4 and S8, the violin plots illustrate how cosine similarity varies as we move from proteins within the same cluster to those in the nearest neighboring clusters and finally to random clusters, highlighting trends for both well-separated and poorly-separated clusters. For all datasets, cosine similarity is notably highest within well-separated clusters, aligning with previous observations on local similarity, while poorly-separated clusters show a more gradual decline. We used Cohen’s D, an effect size measure that quantifies the magnitude of differences between two distributions (values around 0.2 are considered small, ∼ 0.5 medium, and ≥ 0.8 large), to assess two comparisons: (1) intra-cluster versus neighboring-cluster similarity scores, and (2) neighboring-cluster versus random-cluster similarity scores. These comparisons were performed separately for both well-separated and poorly separated clusters. We also report Mann–Whitney U test results (Table S2), which provide a non-parametric measure of statistical significance; together, the two metrics capture both the size and reliability of the observed effects. When comparing proteins to their nearest neighboring cluster, well-separated clusters showed a sharp decrease in similarity relative to within-cluster values, whereas poorly separated clusters showed a more modest decrease (e.g., Cohen’s D = 2.6 vs. D = 0.54, Fig. 4B). This result delineates a clear contrast between regions of the projection where cluster boundaries correspond to embedding-space structure and regions where boundaries are less meaningful. When comparing to a random cluster, the drop in similarity was smaller but more pronounced for poorly separated clusters (e.g., D = 2.0 vs. 0.4, Fig. 4D). This suggests that proteins in poorly separated clusters retain some similarity with nearby clusters in the same cloud. Thus, while absolute distances in 2D should not be overinterpreted, the spatial arrangement of clusters does preserve aspects of the underlying embedding space. While the dimensional reduction serves primarily as a visualization tool, these patterns offer additional context for interpreting both local and global relationships between protein sequences in the visualizations.

In Fig. 4, the violin plots illustrate how cosine similarity varies as we move from proteins within the same cluster to those in the nearest neighboring clusters and finally to random clusters, highlighting trends for both well-separated and poorly-separated clusters. For all three datasets, cosine similarity is notably highest within well-separated clusters, aligning with previous observations on local similarity, while poorly-separated clusters show a more gradual decline. We used Cohen’s D to measure the effect in two comparisons: (1) between intra-cluster similarity scores and neighboring-cluster similarity scores, and (2) between neighboring-cluster similarity scores and random-cluster similarity scores. These comparisons were performed separately for both well-separated and poorly-separated clusters. When measuring similarity with the neighboring cluster, proteins belonging to well-separated clusters show a significant drop in the mean, which is not as noticeable when observing the poorly-separated clusters. On the other hand, similar behavior can be observed as we move farther away from the cluster and measure the similarity of proteins with those in a random cluster, but this time, the proteins situated in a poorly-separated cluster show a more significant drop when compared to proteins in well-separated clusters. This implies that sequences in poorly-separated clusters, located in the “fuzzy”, cloud-like aggregation of clusters, share a higher similarity with their surrounding proteins in the cloud-like formation. This pattern suggests that the spatial relationship in the final representation maintains some meaningful reflection of the underlying data structure, even though the absolute distances should not be interpreted directly. While the dimensional reduction serves primarily as a visualization tool, these patterns offer additional context for interpreting both local and global relationships between protein sequences in the visualizations.

### 2.3 PLVis projections reveal conserved protein families across species

Proteins in organisms have evolved to carry out a wide range of biological functions. As species diverge along the phylogenetic tree, their proteomes shift in content and composition. We reasoned that PLVis projections should be particularly useful in this context, because proteins belonging to conserved families across related species are expected to cluster together in embedding space, making it easier to distinguish broadly conserved groups from lineage-specific proteins. In this section, we focus on two genera of major pathogenic importance, *Mycobacterium* and *Plasmodium*, which include the causative agents of tuberculosis and malaria, respectively.

We first generated a PLM embedding visualization for a subset of species from the genus *Mycobacterium*, a group of over 190 Gram-positive bacterial species belonging to the Actinobacteria phylum. These species range from relatively harmless organisms like *M. smegmatis* to dangerous human pathogens like *M. tuberculosis* and *M. leprae* [45, 46]. These bacteria were traditionally classified by their growth rate (slow or rapid), and recent taxonomic revisions have divided them into five distinct genera: *Mycolicibacterium, Mycolicibacter, Mycolicibacillus, Mycobacteroides*, and *Mycobacterium* [47]. To demonstrate the value that PLVis projections have in comparing proteomes across organisms, we analyzed and visualized the dataset containing the proteomes of eight *Mycobacterium* species: *M. smegmatis, M. fortuitum, M. kansasii, M. marinum, M. leprae, M. tuberculosis, M. bovis*, and *M. intracellulare* (shown in Figure 5).

**Figure 5:**
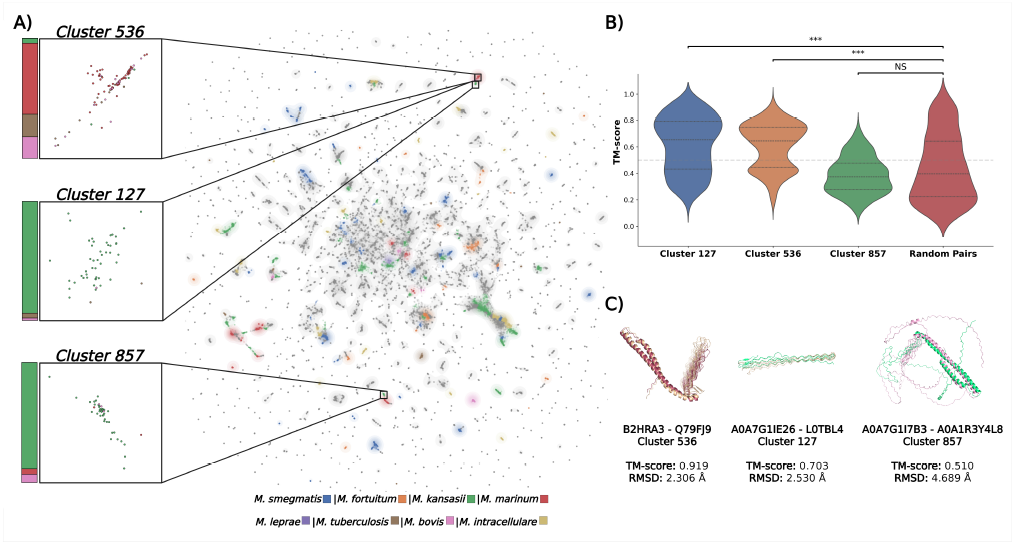
PLVis for the proteomes of eight *Mycobacterium* species and representative protein structures from three clusters. (A) Clusters in the visualization are colored by enrichment for a particular Mycobacterial species (blue: *M. smegmatis*, orange: *M. fortuitum*, green: *M. kansasii*, red: *M. marinum*, purple: *M. leprae*, brown: *M. tuberculosis*, pink: *M. bovis*, yellow: *M. intracellulare*), clusters in gray are not enriched for a Mycobacterium species. The three most enriched clusters in the projection (127, 536 & 857) are zoomed in, and a color bar showing the fraction of organisms in the cluster is located on their left side. (B) Violin plots of the TM-score (measuring structural alignment) between proteins in clusters 127 (blue), 536 (orange), 857 (green), and random pairs (red). Statistical comparison was performed using the Mann-Whitney U test (***p < 0.001) (C) AlphaFold protein structure comparison of proteins in PE-PGRS clusters, colored by organism.

A key insight from visually comparing proteomes across related organisms is the ability to quickly identify which protein families are enriched or expanded in each organism. We thus performed a hypergeometric test with the Benjamini-Hochberg false discovery rate correction to identify the clusters enriched for a single organism. Out of the 1581 k-clusters, 184 ( ∼ 12%) are enriched and are colored according to their respective organisms in Fig. 5. We found that the three clusters with the lowest FDR-corrected p-value (clusters 127, 536 & 857) all contained proteins belonging to the PE-Polymorphic GC-Rich (PE-PGRS) family. These proteins are glycine-rich with multiple GGA/GGN repeats and contain a PE domain near the N-terminus of the sequence as well as a high guanine and cytosine (GC) content of approximately 80% [48, 49, 50]. Cluster 857 contains five glycine-rich “uncharacterized” proteins, one of which (A0A7G1IER6) fulfills all previously mentioned qualities (PE domain, GGX motif, and GC content) of a PE-PGRS family protein. Furthermore, all three clusters were not categorized as well-separated, suggesting that they might be closely related to their neighboring clusters, which is further validated by their positions. Both clusters 127 and 536 are close together and linked with cluster 1389, another enriched cluster with PE-PGRS proteins. Cluster 857, although situated on the other side of the projection, is also surrounded by clusters enriched for PE-PGRS family proteins belonging to *M. marinum* (clusters 149 & 1511). This observation is consistent with recent evolutionary analyses showing that the PE-PGRS family is not a homogeneous group but contains subfamilies and specialized members with distinct roles in mycobacterial pathogenesis and host interaction [51].

To understand why the previously mentioned clusters were positioned in separate parts of the visualization, we obtained the AlphaFold structures for proteins belonging to the PE-PGRS clusters and calculated the TM-score between them. The comparison was made between proteins belonging to the same cluster, and random pairs from different clusters were selected as control, shown in the violin plots of Fig. 5. A close look at the plots for each cluster reveals that the visualization separated the protein family according to similarities in their structure, with clusters 127 and 536 having most of their scores above the 0.5 threshold. Cluster 857 shows lower scores, but a closer look at the structures in the cluster shows that they have a long disordered region near the C-terminus, which could have impacted the structural comparison. Such structural partitioning aligns with reports that individual PE-PGRS proteins have diverged to acquire specialized functions, suggesting that the separation observed here may reflect true biological heterogeneity within this protein family [51]. We thus infer that the projections can separate proteins belonging to the same family according to their structure, which poses a significant advantage when looking for protein analogs to be used in experimental procedures. However, we reiterate that the distance between both groups of clusters is not a measure of their similarity.

Next, we analyzed the *Plasmodium* genus, consisting of protozoan parasites that require a vertebrate and an invertebrate host to complete their life cycle [52]. This genus is medically significant as it contains the parasitic species that cause malaria, a vector-borne infection. Five species within this genus are known to infect humans: *P. falciparum, P. malariae, P. ovale, P. vivax*, and *P. knowlesi* [53]. Similarly to the previous study, we visualized a dataset containing the proteomes of these five parasites, which is shown in Figure 6. Compared to the *Mycobacterium* visualization, the *Plasmodium* PLVis has a larger and central poorly-separated/fuzzy region. (S < 0.5). Of the 1,942 k-clusters, approximately 36% were poorly-separated, compared to 14% in the *Mycobacterium* projection.

**Figure 6:**
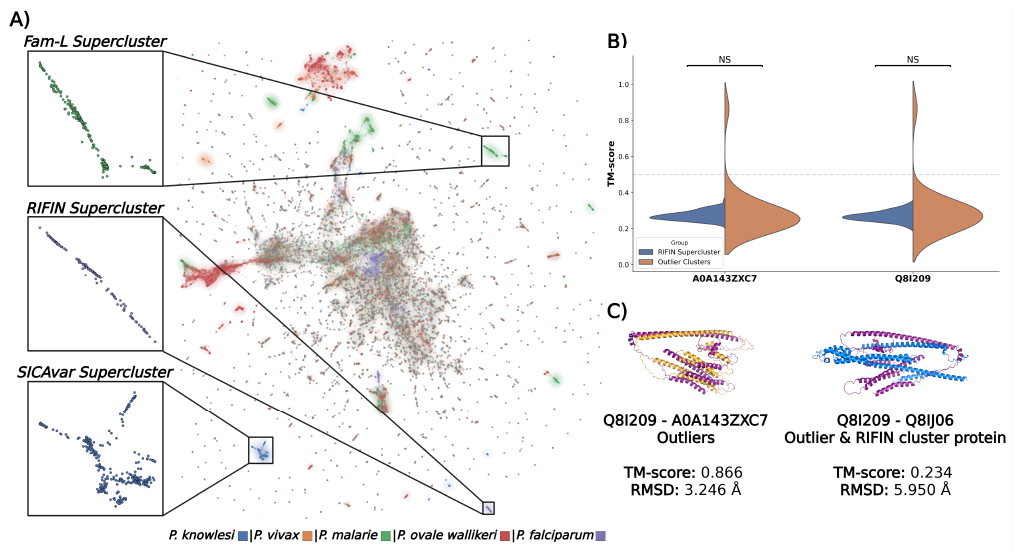
PLVis for the proteomes of five *Plasmodium* species. (A) Proteins in the visualization are colored according to their species (blue: *P. Knowlesi*, orange: *P. vivax*, green: *P. malariae*, red: *P. ovale wallikeri*, purple: *P. falciparum*). The Fam-L, SICAvar, and RIFIN superclusters are zoomed in. (B) Violin plots of the TM-score between outlier proteins (A0A143ZXC7 and Q8I209) with proteins belonging to the RIFIN supercluster (blue) and outlier clusters (orange). (C) AlphaFold protein structure comparison of Q8I209 (outlier) with A0A143ZXC7 (outlier) and Q8IJ06 (RIFIN supercluster).

For this dataset, we repeated the hypergeometric test with the Benjamini-Hochberg false discovery rate correction to identify clusters enriched for a single organism, which resulted in the identification of 375 ( ∼ 19%) enriched clusters. We identified 77 enriched clusters that contained proteins exclusively from a single species, a fact that further exemplifies the greater proteomic diversity of this dataset, due to the more complex organisms shown. Because of this greater diversity, one can quickly point out regions in the projection that highlight a specific family of proteins that belong exclusively to a single species in Figure 6. Such is the case for SICAvar proteins of *P. knowlesi*, Fam-L proteins of *P. malariae*, and RIFIN proteins belonging to *P. falciparum*.

It has been shown that RIFIN proteins are used by *P. falciparum* to evade the host immune system by binding to immune-inhibitory receptors [54]. The visualization reveals that most RIFIN proteins are concentrated in three main clusters (38, 1448, and 1522), with only two RIFIN proteins found elsewhere in clusters 1582 and 1666 (A0A143ZXC7 and Q8I209). These outlier proteins are particularly interesting as they are surrounded by members of multiple protein families (REdfSA, tryptophan-rich antigen (TRAgs), and Maurer’s clefts two transmembrane (PfMC-2TM) proteins) all of which, including RIFIN proteins, are associated with the infected erythrocyte’s membrane [55, 56, 57, 58]. This clustering pattern suggests that A0A143ZXC7 and Q8I209 might also function as erythrocyte surface antigens or membrane proteins. Similarly, the TM-score of these outlier proteins was calculated with the proteins in the RIFIN supercluster and the clusters to which they belong, shown in the violin plots of Fig. 6. We found that the proteins don’t share structural similarity with the supercluster, elucidating why they were positioned apart from the other RIFIN proteins. Yet surprisingly, they also don’t share significant structural similarity with proteins within each cluster. Nonetheless, their embeddings show relatively high similarity to their corresponding cluster proteins (average cosine similarity ∼ 0.7), hinting at a purely functional relation to their neighbors. These observations and associated hypotheses showcase how PLVis can help interactively navigate large-scale protein datasets to reveal biologically significant patterns, while simultaneously providing valuable insights into protein function prediction and pathogen biology.

### 2.4 The PLVis Repository, A Web Portal for Comparative Proteome Analysis Within Taxonomic Families

Having shown the properties of PLVis in the previous sections, we decided to develop an interactive web platform for exploring and comparing UniProtKB reference proteomes, the PLVis Repository. By systematically applying the pipeline across thousands of proteomes and making the results accessible through an interactive interface, the PLVis Repository lowers the barrier for engaging with language model representations, whether in research, teaching, or community-driven discovery. The *Mycobacterium* and *Plasmodium analyses* case studies presented above are likewise integrated into the Repository as examples of its practical use.

Towards capturing comparative full proteome visualizations across the tree of life, we collected all reference proteomes from UniProtKB. This collection of proteomes covers well-studied model organisms and proteomes of interest in biological research [2]. Each proteome comparison is performed at the family level, providing a more balanced distribution that offers taxonomic resolution while including enough species for meaningful comparisons. For each of the available families in UniProtKB, the reference proteomes were retrieved to generate the corresponding PLM embeddings; in the case of outlier taxa with more than ten species, we selected the ten proteomes with the highest BUSCO completeness scores to ensure high-quality and representative comparisons. A total of 4,695 reference proteomes are showcased across 3 domains, 3 kingdoms, 67 phyla, 165 classes, 404 orders, 901 families, and 2,605 genera.

To facilitate navigation across the different taxonomic groups when first accessing the website, users are greeted with a collapsible tree view that helps them explore the available comparisons. Clicking on each taxon expands the tree, showing the next level of available taxa for each rank, showcasing the relation between the comparisons featured in the repository. The website also features search functionality for specific taxonomic ranks, allowing the rapid retrieval of specific proteome visualizations. If a match is found, the page will either present a list of relevant taxonomic families or redirect users to the associated proteome comparison page if the match corresponds to an available family.

Each page contains a list of the available species at the top, separated by “genus”, a PLVis projection of the proteomes, and an enrichment analysis table highlighting overrepresented annotations. Embedding representations for each protein sequence were generated using the ProtT5 language model, followed by dimensionality reduction with the UMAP and t-SNE algorithms. The resulting embeddings were then clustered using K-Means to identify structural and functional patterns within each family. Finally, the K-Means clusters were named by creating bi-grams of the most frequent words in the protein names present in that cluster.

Each visualization is colored according to the organisms shown for their quick identification, as can be seen in Figure 7. A search bar located in the upper left corner of each projection allows users to dynamically filter the data by entering keywords. Users can search by UniProt entry ID, gene name, annotation score, and other metadata fields to highlight specific proteins of interest. By examining the visualization and identifying neighboring proteins in the projection space, users can quickly locate proteins with similar embedding representations that, as demonstrated in previous chapters, preserve meaningful functional and structural information.

**Figure 7:**
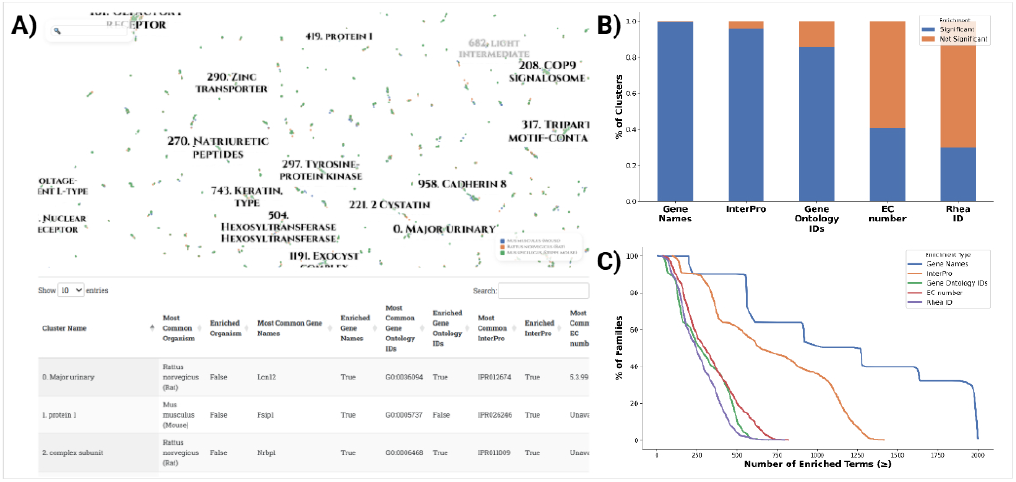
The PLVis Repository shows cluster enrichment of biological significance. (A) Screenshot of the comparison page for the *Muridae* family. The upper half shows the projection with each protein colored by organism. The bottom half shows a table of enriched terms in each cluster. (B) Stacked bar plots showing the proportion of UMAP clusters enriched for genes, InterPro (IP), Gene Ontology (GO), EC numbers, and Rhea IDs. Blue bars indicate the fraction of clusters with statistically significant enrichment (p < 0.05), while orange bars represent clusters without significant enrichment. (C) Cumulative coverage curves depicting the percentage of UMAP clusters with at least N enriched terms, colored by enrichment type.

An enrichment table is located below the visualization to aid users in finding functionally important clusters. Hypergeometric enrichment testing was performed on each cluster by obtaining the most common organism, gene name, InterPro (IP), Gene Ontology (GO), EC number, and Rhea ID, and comparing their distribution within the cluster against all other clusters. This allows users to detect features that are statistically overrepresented and potentially characteristic of specific protein groups. In Figure 7 and S9, we show the percentage of clusters that present functional enrichment across all taxonomic families in the PLVis Repository. More than 80% of UMAP and t-SNE clusters in the repository are enriched for gene, IP, and GO information, with more than 50% of the families having more than 500 significant clusters. However, the proportion of enriched clusters is lower for EC number and Rhea ID when compared to the other features. These trends in enrichment emphasize that the protein clusters available are more strongly aligned with functional domain composition than with metabolic reaction identity. Overall, enrichment analysis demonstrates that the clustering captures biologically meaningful groupings, particularly with respect to protein domains (InterPro) and functional annotations (Gene Ontology).

## 3 Discussion

The PLVis pipeline presented here is an efficient and accessible alternative for the visual representation of protein data obtained from PLM embeddings. When used in conjunction with SSNs, these visualizations enhance protein functional annotation by effectively clustering proteins according to their family classifications. For instance, researchers investigating specific protein families and seeking to validate the function of poorly annotated proteins can utilize PLVis projections to rapidly categorize proteins into distinct subfamilies. This clustering facilitates the identification of promising candidates for experimental validation, particularly when minimally annotated proteins (confidence levels 1 or 2) are found in proximity to well-characterized proteins (confidence level 5).

While the primary strength of PLVis lies in its clustering capabilities, it’s important to understand both its limitations and flexibility in practical applications. As stated before, due to the limitations of dimensionality reduction, distances in the visualizations aren’t meaningful. However, this opens up opportunities for the users to have the liberty to modify cluster coordinates in their datasets, giving meaning to inter-cluster distance based on additional knowledge. For example, clusters can be spatially organized according to various biological parameters, such as gene expression patterns, protein essentiality profiles, or functional categories (e.g., positioning all redox enzymes in a specific region, or separating transcription factors, transporters, and enzymes). This flexibility in visualization emphasizes the importance of domain expertise and underscores the necessity for users to thoroughly understand both their biological data and the analytical tools at their disposal.

Beyond individual protein analysis and cluster organization, PLVis demonstrates remarkable utility in broader comparative studies. From a biological perspective, PLVis projections demonstrate optimal utility in comparative analyses of complete proteomes across different species. The resultant protein clustering patterns reveal significant biological insights, such as species-specific protein family absences or conserved patterns within taxonomic genera. This approach is particularly valuable for analyzing specific biological relationships, exemplified by host-pathogen interactions, where the visualization can identify clusters of proteins from both organisms that may be implicated in pathogenesis. Such protein clusters provide potential molecular signatures associated with disease mechanisms.

The PLVis Repository offers researchers a quick visualization of reference proteome comparisons to find promising protein relationships within taxonomic families. Coupled with the available enrichment tables, the generated projections serve as starting points for deeper biological investigations and hypothesis generation. Expanding the collection of curated case studies through community collaboration could further enhance the website into a better educational and research resource. For this reason, we also make available the link to the PLVis Colab Notebook to assist users in generating their comparisons using the pipeline for the studies above. Together, the PLVis Repository and Colab Notebook provide a scalable platform for visualizing and analyzing large proteomic datasets, helping bridge the gap between massive collections of unannotated proteins and meaningful biological insights.

## References

[1] Valérie de Crécy-lagard, Rocio Amorin de Hegedus, Cecilia Arighi, Jill Babor, Alex Bateman, Ian Blaby, Crysten Blaby-Haas, Alan J Bridge, Stephen K Burley, Stacey Cleveland, Lucy J Colwell, Ana Conesa, Christian Dallago, Antoine Danchin, Anita de Waard, Adam Deutschbauer, Raquel Dias, Yousong Ding, Gang Fang, Iddo Friedberg, John Gerlt, Joshua Goldford, Mark Gorelik, Benjamin M Gyori, Christopher Henry, Geoffrey Hutinet, Marshall Jaroch, Peter D Karp, Liudmyla Kondratova, Zhiyong Lu, Aron Marchler-Bauer, Maria-Jesus Martin, Claire McWhite, Gaurav D Moghe, Paul Monaghan, Anne Morgat, Christopher J Mungall, Darren A Natale, William C Nelson, Seán O’Donoghue, Christine Orengo, Katherine H O’Toole, Predrag Radivojac, Colbie Reed, Richard J Roberts, Dmitri Rodionov, Irina A Rodionova, Jeffrey D Rudolf, Lana Saleh, Gloria Sheynkman, Francoise Thibaud-Nissen, Paul D Thomas, Peter Uetz, David Vallenet, Erica Watson Carter, Peter R Weigele, Valerie Wood, Elisha M Wood-Charlson, and Jin Xu. A roadmap for the functional annotation of protein families: a community perspective. Database, 2022:baac062, January 2022.

[2] The UniProt Consortium. UniProt: the Universal Protein Knowledgebase in 2025. Nucleic Acids Research, 53(D1):D609–D617, January 2025.

[3] Constance J. Jeffery. Current successes and remaining challenges in protein function prediction. Frontiers in Bioinformatics, 3:1222182, July 2023. Publisher: Frontiers.

[4] Kimberly A Reynolds, Eduardo Rosa-Molinar, Robert E Ward, Hongbin Zhang, Breeanna R Urbanowicz, and A Mark Settles. Accelerating Biological Insight for Understudied Genes. Integrative and Comparative Biology, 61(6):2233–2243, December 2021.

[5] Janine N. Copp, Eyal Akiva, Patricia C. Babbitt, and Nobuhiko Tokuriki. Revealing Unexplored Sequence-Function Space Using Sequence Similarity Networks. Biochemistry, 57(31):4651–4662, August 2018. Publisher: American Chemical Society.

[6] Nils Oberg, Timothy W. Precord, Douglas A. Mitchell, and John A. Gerlt. RadicalSAM.org: A Resource to Interpret Sequence-Function Space and Discover New Radical SAM Enzyme Chemistry. ACS Bio & Med Chem Au, 2(1):22–35, February 2022. Publisher: American Chemical Society.

[7] Remi Zallot, Nils Oberg, and John A. Gerlt. Discovery of New Enzymatic Functions and Metabolic Pathways Using Genomic Enzymology Web Tools. Current opinion in biotechnology, 69:77, January 2021.

[8] Amra Dhabalia Ashok, Jella N. Freitag, Iker Irisarri, Sophie de Vries, and Jan de Vries. Sequence similarity networks bear out hierarchical relationships of green cytochrome P450. Physiologia plantarum, 176(2):e14244, 2024.

[9] Audrey R. Long, Emma L. Mortara, Brisa N. Mendoza, Emma C. Fink, Francis X. Sacco, Matthew J. Ciesla, and Tyler M. M. Stack. Sequence similarity network analysis of drug- and dye-modifying azoreductase enzymes found in the human gut microbiome. Archives of Biochemistry and Biophysics, 757:110025, July 2024.

[10] Holly J. Atkinson, John H. Morris, Thomas E. Ferrin, and Patricia C. Babbitt. Using Sequence Similarity Networks for Visualization of Relationships Across Diverse Protein Superfamilies. PLoS ONE, 4(2):e4345, February 2009.

[11] Tancred Frickey and Andrei Lupas. CLANS: a Java application for visualizing protein families based on pairwise similarity. Bioinformatics, 20(18):3702–3704, December 2004.

[12] John A. Gerlt, Jason T. Bouvier, Daniel B. Davidson, Heidi J. Imker, Boris Sadkhin, David R. Slater, and Katie L. Whalen. Enzyme Function Initiative-Enzyme Similarity Tool (EFI-EST): A web tool for generating protein sequence similarity networks. Biochimica et biophysica acta, 1854(8):1019, April 2015.

[13] SR Eddy. Profile hidden Markov models. Bioinformatics, 14(9):755–763, January 1998.

[14] Simon C. Potter, Aurélien Luciani, Sean R. Eddy, Youngmi Park, Rodrigo Lopez, and Robert D. Finn. HMMER web server: 2018 update. Nucleic Acids Research, 46(Web Server issue):W200, June 2018.

[15] L.R. Rabiner. A tutorial on hidden Markov models and selected applications in speech recognition. Proceedings of the IEEE, 77(2):257–286, February 1989. Conference Name: Proceedings of the IEEE.

[16] Bhavya Mor, Sunita Garhwal, and Ajay Kumar. A Systematic Review of Hidden Markov Models and Their Applications. Archives of Computational Methods in Engineering, 28(3):1429–1448, May 2021.

[17] Benjamin Schuster-Böckler, Jörg Schultz, and Sven Rahmann. HMM Logos for visualization of protein families. BMC Bioinformatics, 5(1):7, January 2004.

[18] Travis J. Wheeler, Jody Clements, and Robert D. Finn. Skylign: a tool for creating informative, interactive logos representing sequence alignments and profile hidden Markov models. BMC Bioinformatics, 15(1):7, January 2014.

[19] Adam Krejci, Ted R. Hupp, Matej Lexa, Borivoj Vojtesek, and Petr Muller. Hammock: a hidden Markov model-based peptide clustering algorithm to identify protein-interaction consensus motifs in large datasets. Bioinformatics, 32(1):9–16, January 2016.

[20] Inigo Barrio-Hernandez, Jingi Yeo, Jürgen Jänes, Milot Mirdita, Cameron L. M. Gilchrist, Tanita Wein, Mihaly Varadi, Sameer Velankar, Pedro Beltrao, and Martin Steinegger. Clustering predicted structures at the scale of the known protein universe. Nature, 622(7983):637–645, 2023.

[21] Dan Ofer, Nadav Brandes, and Michal Linial. The language of proteins: NLP, machine learning & protein sequences. Computational and Structural Biotechnology Journal, 19:1750, March 2021.

[22] Tristan Bepler and Bonnie Berger. Learning the Protein Language: Evolution, Structure and Function. Cell systems, 12(6):654, June 2021.

[23] Ali Madani, Ben Krause, Eric R. Greene, Subu Subramanian, Benjamin P. Mohr, James M. Holton, Jose Luis Olmos, Caiming Xiong, Zachary Z. Sun, Richard Socher, James S. Fraser, and Nikhil Naik. Large language models generate functional protein sequences across diverse families. Nature Biotechnology, 41(8):1099–1106, August 2023. Publisher: Nature Publishing Group.

[24] Abel Chandra, Laura Tünnermann, Tommy Löfstedt, and Regina Gratz. Transformer-based deep learning for predicting protein properties in the life sciences. eLife, 12:e82819, January 2023. Publisher: eLife Sciences Publications, Ltd.

[25] Noelia Ferruz, Steffen Schmidt, and Birte Höcker. ProtGPT2 is a deep unsupervised language model for protein design. Nature Communications, 13:4348, July 2022.

[26] Baohui Lin, Xiaoling Luo, Yumeng Liu, and Xiaopeng Jin. A comprehensive review and comparison of existing computational methods for protein function prediction. Briefings in Bioinformatics, 25(4):bbae289, July 2024.

[27] Robert Schmirler, Michael Heinzinger, and Burkhard Rost. Fine-tuning protein language models boosts predictions across diverse tasks. Nature Communications, 15(1):7407, August 2024. Publisher: Nature Publishing Group.

[28] Michael Heinzinger, Maria Littmann, Ian Sillitoe, Nicola Bordin, Christine Orengo, and Burkhard Rost. Contrastive learning on protein embeddings enlightens midnight zone. NAR Genomics and Bioinformatics, 4(2):lqac043, June 2022.

[29] Wayland Yeung, Zhongliang Zhou, Liju Mathew, Nathan Gravel, Rahil Taujale, Brady O’Boyle, Mariah Salcedo, Aarya Venkat, William Lanzilotta, Sheng Li, and Natarajan Kannan. Tree visualizations of protein sequence embedding space enable improved functional clustering of diverse protein superfamilies. Briefings in Bioinformatics, 24(1):bbac619, January 2023.

[30] Tobias Senoner, Tobias Olenyi, Michael Heinzinger, Anton Spannagl, George Bouras, Burkhard Rost, and Ivan Koludarov. ProtSpace: A Tool for Visualizing Protein Space. Journal of Molecular Biology, 437(15):168940, August 2025.

[31] Ami G. Sangster, Cameron Dufault, Haoning Qu, Denise Le, Julie D. Forman-Kay, and Alan M. Moses. Zeroshot segmentation using embeddings from a protein language model identifies functional regions in the human proteome. PLOS Computational Biology, 21(11):e1012929, November 2025. Publisher: Public Library of Science.

[32] Tara Chari and Lior Pachter. The specious art of single-cell genomics. PLOS Computational Biology, 19(8):e1011288, August 2023. Publisher: Public Library of Science.

[33] Dmitry Kobak and George C. Linderman. Initialization is critical for preserving global data structure in both t-SNE and UMAP. Nature Biotechnology, 39(2):156–157, February 2021. Publisher: Nature Publishing Group.

[34] Alex Diaz-Papkovich, Luke Anderson-Trocmé, Chief Ben-Eghan, and Simon Gravel. UMAP reveals cryptic population structure and phenotype heterogeneity in large genomic cohorts. PLOS Genetics, 15(11):e1008432, November 2019. Publisher: Public Library of Science.

[35] Anant Dadu, Vipul K. Satone, Rachneet Kaur, Mathew J. Koretsky, Hirotaka Iwaki, Yue A. Qi, Daniel M. Ramos, Brian Avants, Jacob Hesterman, Roger Gunn, Mark R. Cookson, Michael E. Ward, Andrew B. Singleton, Roy H. Campbell, Mike A. Nalls, and Faraz Faghri. Application of Aligned-UMAP to longitudinal biomedical studies. Patterns, 4(6):100741, June 2023.

[36] Shu Wang, Eduardo D. Sontag, and Douglas A. Lauffenburger. What Cannot Be Seen Correctly in 2D Visualizations Of Single-Cell ‘Omics Data? Cell systems, 14(9):723, September 2023.

[37] Ahmed Elnaggar, Michael Heinzinger, Christian Dallago, Ghalia Rehawi, Yu Wang, Llion Jones, Tom Gibbs, Tamas Feher, Christoph Angerer, Martin Steinegger, Debsindhu Bhowmik, and Burkhard Rost. ProtTrans: Toward Understanding the Language of Life Through Self-Supervised Learning. IEEE Transactions on Pattern Analysis and Machine Intelligence, 44(10):7112–7127, October 2022. Conference Name: IEEE Transactions on Pattern Analysis and Machine Intelligence.

[38] Thomas Hayes, Roshan Rao, Halil Akin, Nicholas J. Sofroniew, Deniz Oktay, Zeming Lin, Robert Verkuil, Vincent Q. Tran, Jonathan Deaton, Marius Wiggert, Rohil Badkundri, Irhum Shafkat, Jun Gong, Alexander Derry, Raul S. Molina, Neil Thomas, Yousuf A. Khan, Chetan Mishra, Carolyn Kim, Liam J. Bartie, Matthew Nemeth, Patrick D. Hsu, Tom Sercu, Salvatore Candido, and Alexander Rives. Simulating 500 million years of evolution with a language model. Science, 387(6736):850–858, February 2025. Publisher: American Association for the Advancement of Science.

[39] Leland McInnes, John Healy, Nathaniel Saul, and Lukas Großberger. UMAP: Uniform Manifold Approximation and Projection. Journal of Open Source Software, 3(29):861, September 2018.

[40] Ehsan Amid and Manfred K. Warmuth. TriMap: Large-scale Dimensionality Reduction Using Triplets, March 2022. 1910.00204.

[41] Yingfan Wang, Haiyang Huang, Cynthia Rudin, and Yaron Shaposhnik. Understanding how dimension reduction tools work: an empirical approach to deciphering t-SNE, UMAP, TriMap, and PaCMAP for data visualization. J. Mach. Learn. Res., 22(1):201:9129–201:9201, January 2021.

[42] Suwen Zhao, Ayano Sakai, Xinshuai Zhang, Matthew W Vetting, Ritesh Kumar, Brandan Hillerich, Brian San Francisco, Jose Solbiati, Adam Steves, Shoshana Brown, Eyal Akiva, Alan Barber, Ronald D Seidel, Patricia C Babbitt, Steven C Almo, John A Gerlt, and Matthew P Jacobson. Prediction and characterization of enzymatic activities guided by sequence similarity and genome neighborhood networks. eLife, 3:e03275, June 2014. Publisher: eLife Sciences Publications, Ltd.

[43] Katherine H. O’Toole, Barbara Imperiali, and Karen N. Allen. Glycoconjugate pathway connections revealed by sequence similarity network analysis of the monotopic phosphoglycosyl transferases. Proceedings of the National Academy of Sciences of the United States of America, 118(4):e2018289118, January 2021.

[44] Angela Giorgianni, Alice Zenone, Leander Sützl, Florian Csarman, and Roland Ludwig. Exploring class III cellobiose dehydrogenase: sequence analysis and optimized recombinant expression. Microbial Cell Factories, 23(1):146, May 2024.

[45] Enrico Tortoli, Tarcisio Fedrizzi, Conor J. Meehan, Alberto Trovato, Antonella Grottola, Elisabetta Giacobazzi, Giulia Fregni Serpini, Sara Tagliazucchi, Anna Fabio, Clotilde Bettua, Roberto Bertorelli, Francesca Frascaro, Veronica De Sanctis, Monica Pecorari, Olivier Jousson, Nicola Segata, and Daniela M. Cirillo. The new phylogeny of the genus Mycobacterium: The old and the news. Infection, Genetics and Evolution: Journal of Molecular Epidemiology and Evolutionary Genetics in Infectious Diseases, 56:19–25, December 2017.

[46] Nathan L. Bachmann, Rauf Salamzade, Abigail L. Manson, Richard Whittington, Vitali Sintchenko, Ashlee M. Earl, and Ben J. Marais. Key Transitions in the Evolution of Rapid and Slow Growing Mycobacteria Identified by Comparative Genomics. Frontiers in Microbiology, 10:3019, January 2020. Publisher: Frontiers.

[47] Radhey S. Gupta, Brian Lo, and Jeen Son. Phylogenomics and Comparative Genomic Studies Robustly Support Division of the Genus Mycobacterium into an Emended Genus Mycobacterium and Four Novel Genera. Frontiers in Microbiology, 9:67, February 2018. Publisher: Frontiers.

[48] Flavio De Maio, Rita Berisio, Riccardo Manganelli, and Giovanni Delogu. PE_pgrs proteins of Mycobacterium tuberculosis: A specialized molecular task force at the forefront of host-pathogen interaction. Virulence, 11(1):898– 915, December 2020.

[49] Christopher D’Souza, Uday Kishore, and Anthony G. Tsolaki. The PE-PPE Family of Mycobacterium tuberculosis: Proteins in Disguise. Immunobiology, 228(2):152321, March 2023.

[50] Eliza Kramarska, Flavio De Maio, Giovanni Delogu, and Rita Berisio. Structural Basis of PE_pgrs Polymorphism, a Tool for Functional Modulation. Biomolecules, 13(5):812, May 2023.

[51] Bei Chen, Belmin Bajramović, Bastienne Vriesendorp, and Herman Pieter Spaink. Evolution of the PE_pgrs Proteins of Mycobacteria: Are All Equal or Are Some More Equal than Others? Biology, 14(3):247, February 2025.

[52] IW Sherman. Biochemistry of Plasmodium (malarial parasites). Microbiological Reviews, 43(4):453–495, December 1979. Publisher: American Society for Microbiology.

[53] Spinello Antinori, Laura Galimberti, Laura Milazzo, and Mario Corbellino. Biology of human malaria plasmodia including Plasmodium knowlesi. Mediterranean Journal of Hematology and Infectious Diseases, 4(1):e2012013, 2012.

[54] Fumiji Saito, Kouyuki Hirayasu, Takeshi Satoh, Christian W. Wang, John Lusingu, Takao Arimori, Kyoko Shida, Nirianne Marie Q. Palacpac, Sawako Itagaki, Shiroh Iwanaga, Eizo Takashima, Takafumi Tsuboi, Masako Kohyama, Tadahiro Suenaga, Marco Colonna, Junichi Takagi, Thomas Lavstsen, Toshihiro Horii, and Hisashi Arase. Immune evasion of Plasmodium falciparum by RIFIN via inhibitory receptors. Nature, 552(7683):101–105, December 2017. Publisher: Nature Publishing Group.

[55] A. F. Cowman, R. L. Coppel, R. B. Saint, J. Favaloro, P. E. Crewther, H. D. Stahl, A. E. Bianco, G. V. Brown, R. F. Anders, and D. J. Kemp. The ring-infected erythrocyte surface antigen (RESA) polypeptide of Plasmodium falciparum contains two separate blocks of tandem repeats encoding antigenic epitopes that are naturally immunogenic in man. Molecular Biology & Medicine, 2(3):207–221, June 1984.

[56] Bo Wang, Feng Lu, Yang Cheng, Jun-Hu Chen, Hye-Yoon Jeon, Kwon-Soo Ha, Jun Cao, Myat Htut Nyunt, Jin-Hee Han, Seong-Kyun Lee, Myat Phone Kyaw, Jetsumon Sattabongkot, Eizo Takashima, Takafumi Tsuboi, and Eun-Taek Han. Immunoprofiling of the Tryptophan-Rich Antigen Family in Plasmodium vivax. Infection and Immunity, 83(8):3083–3095, July 2015. Publisher: American Society for Microbiology.

[57] Iryna Tsarukyanova, Judy A. Drazba, Hisashi Fujioka, Satya P. Yadav, and Tobili Y. Sam-Yellowe. Proteins of the Plasmodium falciparum two transmembrane Maurer’s cleft protein family, PfMC-2TM, and the 130 kDa Maurer’s cleft protein define different domains of the infected erythrocyte intramembranous network. Parasitology Research, 104(4):875–891, March 2009.

[58] Mohamed S. Abdel-Latif, Klaus Dietz, Saadou Issifou, Peter G. Kremsner, and Mo-Quen Klinkert. Antibodies to Plasmodium falciparum rifin proteins are associated with rapid parasite clearance and asymptomatic infections. Infection and Immunity, 71(11):6229–6233, November 2003.

