## Supplemental Information for "Protein Language Visualizer: A Repository for Homology Exploration with Language Model Embeddings"

[1]*Department of Chemistry and Biochemistry, University of California San Diego, 9500 Gilman Dr., CA 92093, United States*
[2]*Unidad Profesional Interdisciplinaria de Ingeniería Campus Guanajuato IPN, Instituto Politécnico Nacional, Av. Mineral de Valenciana No. 200, GTO 36275, Mexico*
[3]*Facultad de Medicina Mexicali, Universidad Autónoma de Baja California, Dr. Humberto Torres Sanginés S/N, BC 21000, Mexico*
[4]*Escuela Superior de Medicina, Instituto Politécnico Nacional, Plan de San Luis y Díaz Mirón S/N, CDMX 11340, Mexico*
[5]*Instituto Tecnológico y de Estudios Superiores de Occidente, Anillo Perif. Sur Manuel Gómez Morín 8585, JAL 45604, Mexico.*
[6]*Skaggs School of Pharmacy and Pharmaceutical Sciences, University of California San Diego, 9500 Gilman Dr., CA 92093, United States*

### 1 Sources and Reproducibility

The results and the PLVis Repository shown in this manuscript were developed using Python 3.12 and Jupyter Notebook [1]. Data processing and calculations were obtained using the following Python libraries: numpy (ver.2.1), pandas (ver.2.2), Scikit-Learn (ver.1.6), Scipy (ver.1.15), trimap (ver.1.1), umap-learn (ver.0.5) [2, 3, 4, 5, 6, 7]. Datamapplot (ver.0.5), Matplotlib (ver.3.10), and Seaborn (ver.0.13) were used to generate the plots in the manuscript [8, 9].

All of the information in the datasets was retrieved from the UniProt database (release 2024_06) [10]. All of the structural representations used in our findings were retrieved from the AlphaFold database v4 [11]. Not all sequences contained the same level of annotation in UniProt. Several entries lacked metadata such as EC numbers, Rhea ID, Gene Ontology ID, and InterPro ID. These omissions reflect the available information in UniProt at the time of curation rather than filtering on our part. Likewise, not all proteins had available predicted structures in the AlphaFold Database at the time of curation. No additional annotation or imputation was performed, and all analyses were carried out using the available information for each entry.

#### 2 Supplemental Tables

**Table S1: Summary of UMAP-PLVis protein dataset properties.**

| Dataset | Sequences | Clusters | Well separated clusters* | Poorly separated clusters† |
|---|---|---|---|---|
| *M. tuberculosis* | 3,995 | 96 | 9 | 49 |
| *Mycobacterium* | 41,741 | 1,581 | 667 | 229 |
| *Plasmodium* | 48,647 | 1,942 | 568 | 701 |
| rSAM | 10,000 | 88 | 11 | 19 |
| Sterol proteins | 5,598 | 48 | 8 | 6 |

* Protein sequences with a UniProt annotation score < 3

† Clusters with an average silhouette score >= 0.95

**Table S2: Mann–Whitney U test results for comparisons of within-cluster (CS), nearest-neighbor (NNS), and random-neighbor (RNS) similarities for well and poorly-separated clusters.**

| Dataset | Cluster type | CS-NNS U* | CS-NNS p* | NNS-RNS U† | NNS-RNS p† |
|---|---|---|---|---|---|
| *M. tuberculosis* | Well-separated | 3.389e7 | < 1e-308 | 5.601e7 | 6.862e-274 |
| | Poorly-separated | 4.463e9 | < 1e-308 | 9.232e9 | < 1e-308 |
| *Mycobacterium* | Well-separated | 8.909e9 | < 1e-308 | 1.605e10 | < 1e-308 |
| | Poorly-separated | 6.845e10 | < 1e-308 | 9.999e10 | < 1e-308 |
| *Plasmodium* | Well-separated | 4.757e9 | < 1e-308 | 7.933e9 | < 1e-308 |
| | Poorly-separated | 2.211e11 | < 1e-308 | 4.773e11 | < 1e-308 |
| rSAM | Well-separated | 1.360e8 | < 1e-308 | 6.955e8 | < 1e-308 |
| | Poorly-separated | 6.643e10 | < 1e-308 | 1.065e11 | < 1e-308 |
| Sterol proteins | Well-separated | 3.025e7 | < 1e-308 | 2.070e8 | 2.049e-1 |
| | Poorly-separated | 4.523e9 | < 1e-308 | 1.035e10 | < 1e-308 |

* Same cluster and nearest neighbor comparison values.

† Nearest neighbor and random neighbor comparison values.

### 3 Supplemental Figures

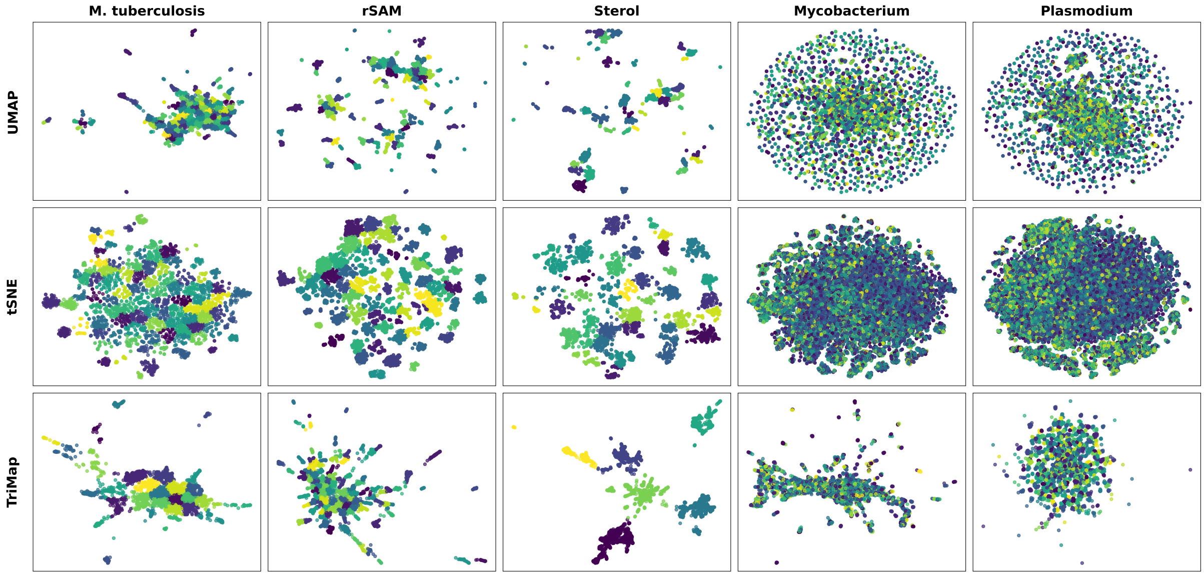


**Figure S1: Dimensionality reduction of protein datasets using UMAP, t-SNE, and TriMap. Projections of *M. tuberculosis* proteome, rSAM proteins, sterol-binding proteins, eight *Mycobacterium* species proteome, and five *Plasmodium* species proteome were generated.**

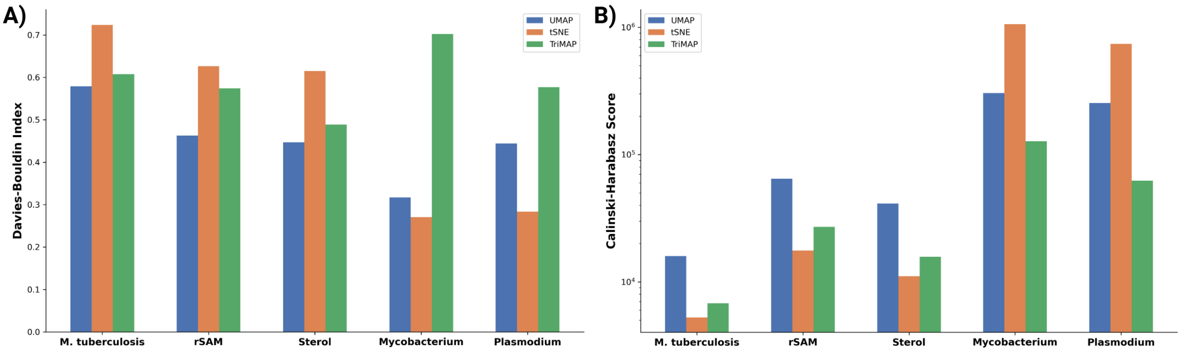


**Figure S2: Comparison of clustering quality across five protein datasets (*M. tuberculosis* proteome, rSAM proteins, sterol-binding proteins, eight *Mycobacterium* species proteome, and five *Plasmodium* species proteome). (A) Davies-Bouldin indices and (B) Calinski-Harabasz scores. UMAP -blue, tSNE - orange, TriMap - green.**

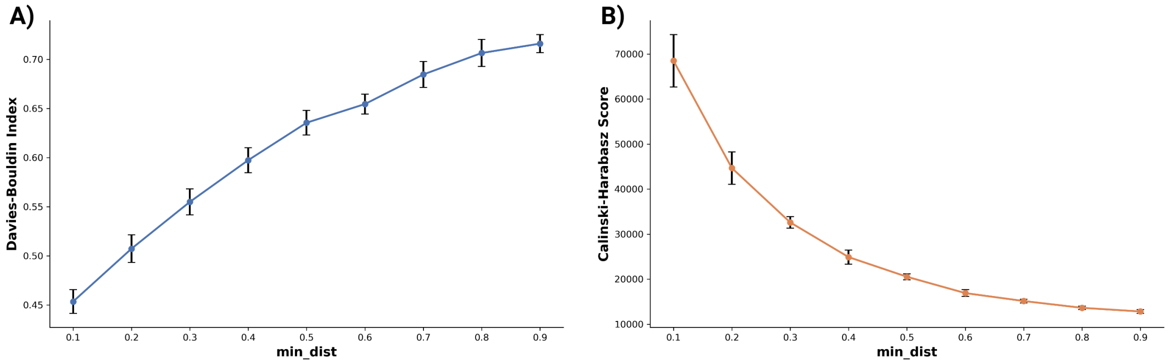


**Figure S3: Effect of the UMAP min_dist hyperparameter on cluster quality. (A) Davies-Bouldin Index and (B) Calinski-Harabasz Index were computed after dimensionality reduction with UMAP followed by K-means clustering. Each point shows the mean metric value across 10 random seeds, and error bars represent the standard deviation.**

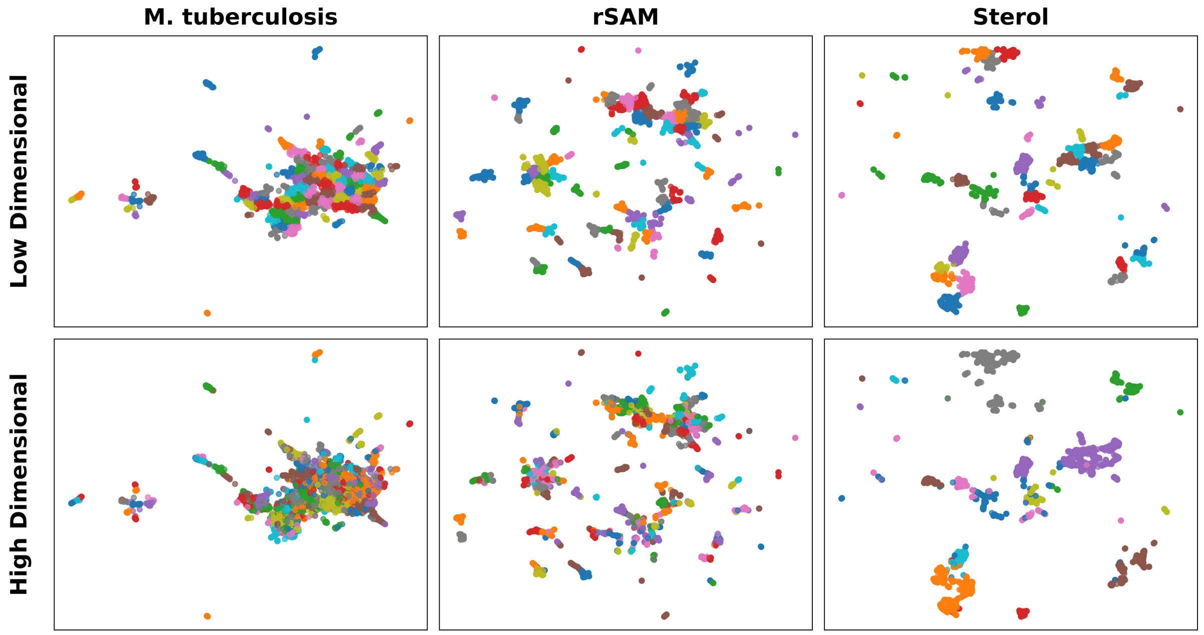


**Figure S4: Comparison of K-means clustering in low-dimensional (top) and high-dimensional (bottom) embeddings across three protein datasets (*M. tuberculosis* proteome, rSAM proteins, sterol-binding proteins). Each point represents a protein and is colored by its K-means cluster assignment. Clustering in the two-dimensional projection yields visually well-separated groups that align with the structure shown in the map, facilitating interpretability of the visualization. In contrast, when the same clustering procedure is applied in the original high-dimensional embedding space, cluster boundaries are less visually separable when projected to two dimensions.**

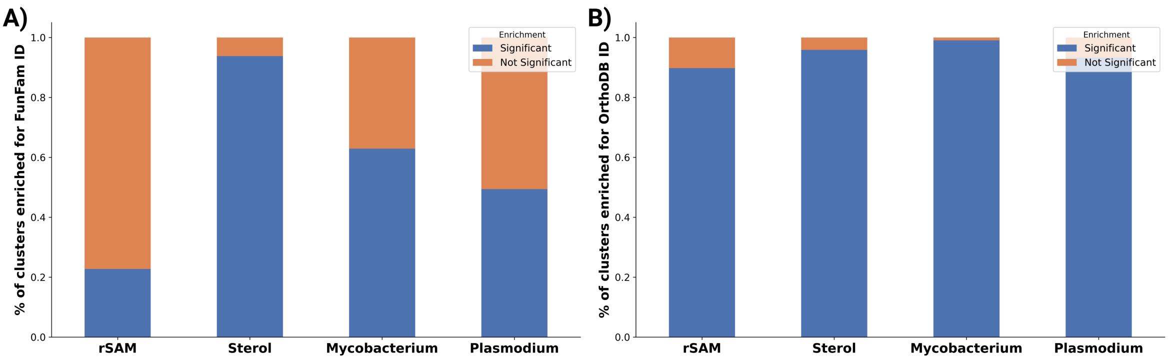


**Figure S5: Stacked bar plots showing the proportion of K-means clusters enriched for at least one CATH FunFam or OrthoDB ID. Enrichment was assessed for CATH FunFams (A) and OrthoDB ortholog groups (B) across the rSAM, sterol, *Mycobacterium*, and *Plasmodium* datasets. Blue bars indicate the fraction of clusters with statistically significant enrichment (adjusted $p < 0.05$), while orange bars represent clusters without significant enrichment. Differences in enrichment levels between FunFams and OrthoDB reflect differences in annotation coverage and scope, with OrthoDB providing broader coverage across species, whereas FunFam annotations are more limited for large and diverse enzyme families.**

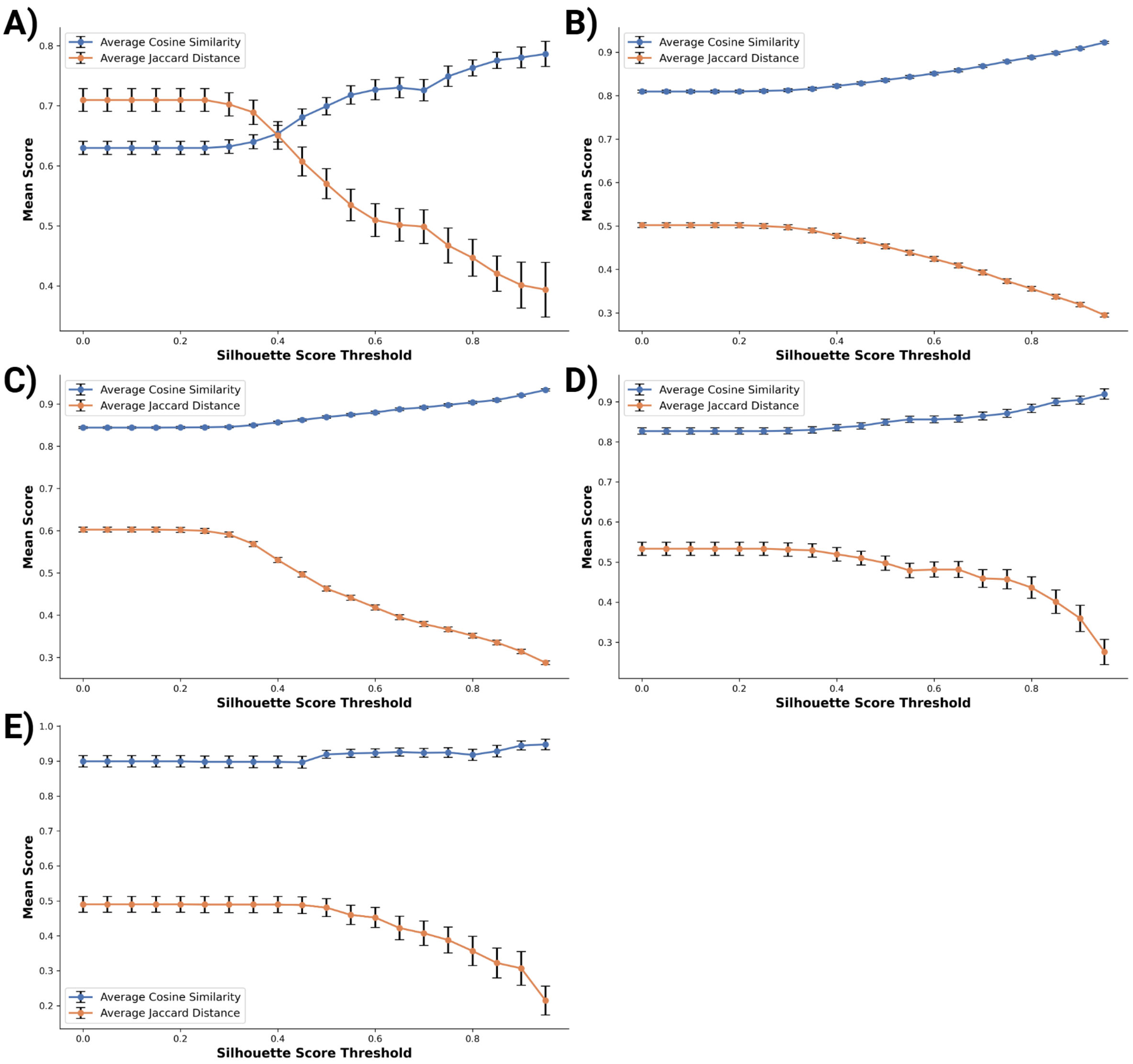


**Figure S6: Relationship between cluster quality and intra-cluster similarity across datasets.** (A) Full proteome of *M. tuberculosis*, (B) eight *Mycobacterium* proteomes, (C) five *Plasmodium* proteomes, (D) 10,000 radical SAM (rSAM) enzymes, and (E) set of sterol-binding proteins. For each, clusters were stratified by their mean silhouette score, and only those above the indicated threshold were retained. The mean intra-cluster cosine similarity (blue) and Jaccard distance (orange) were then computed for the selected clusters. Points represent mean values across clusters, and error bars indicate the standard error of the mean.

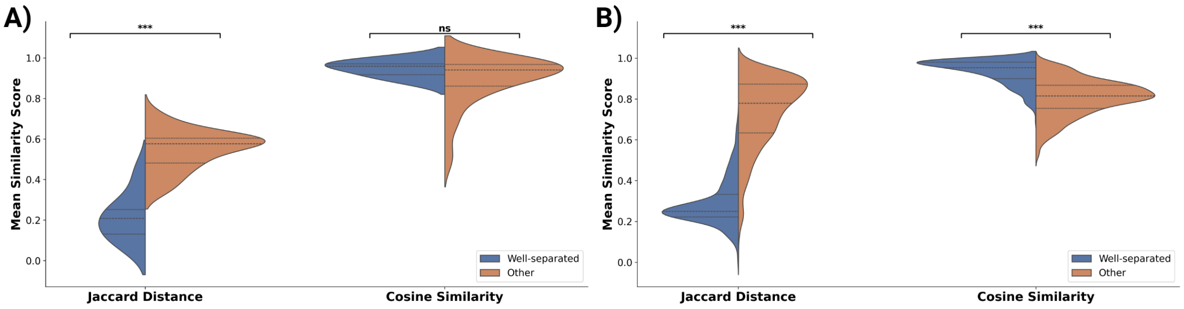


**Figure S7:** Violin plots of the average Jaccard distance of proteins and cosine similarity of high-dimensional embeddings within well-separated clusters (blue) and other clusters (orange) for (A) sterol protein dataset, (B) *Plasmodium* genus proteomes; blue - well-separated clusters, detected by silhouette score above threshold ($S \geq 0.95$), orange - clusters with a silhouette score below the threshold.

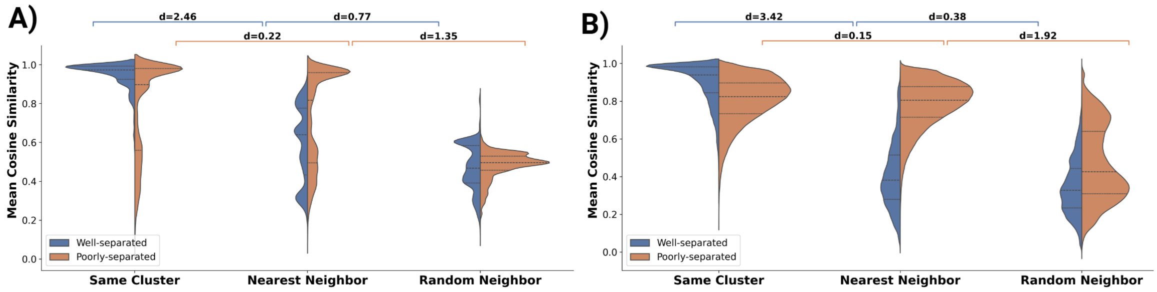


**Figure S8:** Cosine similarity score violin plots for each cluster when comparing its proteins with the nearest neighboring cluster and a randomly selected cluster for (A) sterol protein dataset, (B) *Plasmodium* genus proteomes; blue - well-separated clusters, detected by silhouette score above threshold ($S \geq 0.95$), orange - poorly-separated clusters ($S < 0.5$).

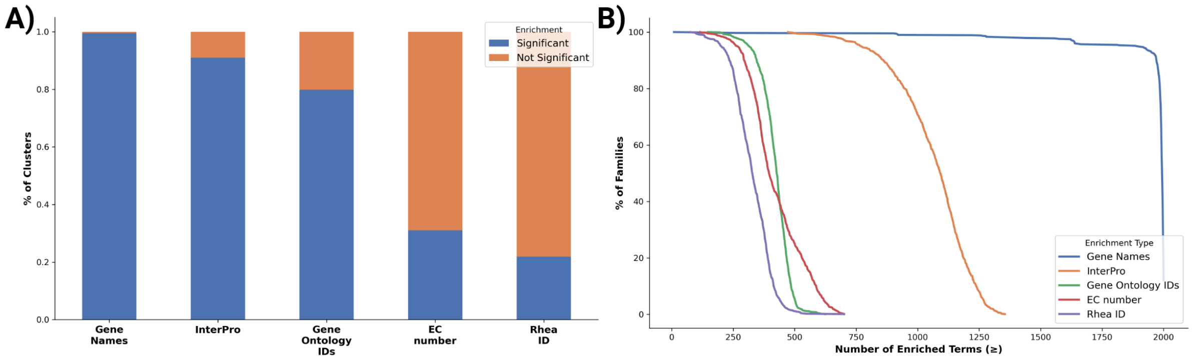


**Figure S9:** Stacked bar plots showing the proportion of t-SNE clusters enriched for genes, InterPro (IP), Gene Ontology (GO), EC numbers, and Rhea IDs. Blue bars indicate the fraction of clusters with statistically significant enrichment (adjusted $p < 0.05$), while orange bars represent clusters without significant enrichment. (C) Cumulative coverage curves depicting the percentage of t-SNE clusters with at least N enriched terms, colored by enrichment type.

### References

[1] Brian E. Granger and Fernando Pérez. Jupyter: Thinking and Storytelling With Code and Data. *Computing in Science & Engineering*, 23(2):7–14, March 2021.

[2] Charles R. Harris, K. Jarrod Millman, Stéfan J. van der Walt, Ralf Gommers, Pauli Virtanen, David Cournapeau, Eric Wieser, Julian Taylor, Sebastian Berg, Nathaniel J. Smith, Robert Kern, Matti Picus, Stephan Hoyer, Marten H. van Kerkwijk, Matthew Brett, Allan Haldane, Jaime Fernández del Río, Mark Wiebe, Pearu Peterson, Pierre Gérard-Marchant, Kevin Sheppard, Tyler Reddy, Warren Weckesser, Hameer Abbasi, Christoph Gohlke, and Travis E. Oliphant. Array programming with NumPy. *Nature*, 585(7825):357–362, September 2020. Publisher: Nature Publishing Group.

[3] Wes McKinney. Data Structures for Statistical Computing in Python. *SciPy 2010*, May 2010.

[4] Fabian Pedregosa, Gaël Varoquaux, Alexandre Gramfort, Vincent Michel, Bertrand Thirion, Olivier Grisel, Mathieu Blondel, Peter Prettenhofer, Ron Weiss, Vincent Dubourg, Jake Vanderplas, Alexandre Passos, David Cournapeau, Matthieu Brucher, Matthieu Perrot, and Édouard Duchesnay. Scikit-learn: Machine Learning in Python. *Journal of Machine Learning Research*, 12(85):2825–2830, 2011.

[5] Pauli Virtanen, Ralf Gommers, Travis E. Oliphant, Matt Haberland, Tyler Reddy, David Cournapeau, Evgeni Burovski, Pearu Peterson, Warren Weckesser, Jonathan Bright, Stéfan J. van der Walt, Matthew Brett, Joshua Wilson, K. Jarrod Millman, Nikolay Mayorov, Andrew R. J. Nelson, Eric Jones, Robert Kern, Eric Larson, C. J. Carey, İlhan Polat, Yu Feng, Eric W. Moore, Jake VanderPlas, Denis Laxalde, Josef Perktold, Robert Cimrman, Ian Henriksen, E. A. Quintero, Charles R. Harris, Anne M. Archibald, Antônio H. Ribeiro, Fabian Pedregosa, and Paul van Mulbregt. SciPy 1.0: fundamental algorithms for scientific computing in Python. *Nature Methods*, 17(3):261–272, March 2020. Publisher: Nature Publishing Group.

[6] Ehsan Amid and Manfred K. Warmuth. TriMap: Large-scale Dimensionality Reduction Using Triplets, March 2022. arXiv:1910.00204 [cs].

[7] Leland McInnes, John Healy, Nathaniel Saul, and Lukas Großberger. UMAP: Uniform Manifold Approximation and Projection. *Journal of Open Source Software*, 3(29):861, September 2018.

[8] John D. Hunter. Matplotlib: A 2D Graphics Environment. *Computing in Science & Engineering*, 9(3):90–95, May 2007.

[9] Michael L. Waskom. seaborn: statistical data visualization. *Journal of Open Source Software*, 6(60):3021, April 2021.

[10] The UniProt Consortium. UniProt: the Universal Protein Knowledgebase in 2025. *Nucleic Acids Research*, 53(D1):D609–D617, January 2025.

[11] Mihaly Varadi, Damian Bertoni, Paulyna Magana, Urmila Paramval, Ivanna Pidruchna, Malarvizhi Radhakrishnan, Maxim Tsenkov, Sreenath Nair, Milot Mirdita, Jingi Yeo, Oleg Kovalevskiy, Kathryn Tunyasuvunakool, Agata Laydon,

Augustin Žídek, Hamish Tomlinson, Dhavanthi Hariharan, Josh Abrahamson, Tim Green, John Jumper, Ewan Birney, Martin Steinegger, Demis Hassabis, and Sameer Velankar. AlphaFold Protein Structure Database in 2024: providing structure coverage for over 214 million protein sequences. *Nucleic Acids Research*, 52(D1):D368–D375, January 2024.
